# Can Harvesting Flowers Reduce the Amount of Atmospheric Carbon Dioxide? : The Case of Cherry Blossoms

**DOI:** 10.1101/2022.10.26.513945

**Authors:** Hannah Zo, Stephen J. Appleyard

## Abstract

Cherry blossoms are popular as street trees in East Asia, providing an attractive backdrop to urban architecture, however their fallen petals can create a waste problem. These petals are likely to contain a significant proportion of fixed carbon so it is suggested that harvesting them may be a solution to both the waste problem and a means of quenching atmospheric CO_2_ concomitantly. This study investigated the feasibility of flower harvesting for reducing atmospheric CO_2_. In particular, the total carbon (TC) stored in all cherry blossoms on streets was quantified in the geographic area of South Korea and compared to various CO_2_ emission rates or amounts quenched by other methods. Branches with flowers were collected from different locations; the TC stored in them ranged between 41.5% - 44.8% of flower dry weights, resulting a mean flower TC per a metre of branch as 0.851±0.070 gC/m. A functional relationship of the sum of the two most apical branch lengths against crown diameter was developed to obtain an estimate of total flowering branch length from the crown diameter of a typical tree on street. The product of flowering branch length and flower TC per a metre of branch indicated that TC stored in all flowers of a tree summed to 336±163 g of carbon, equivalent to 1.23±0.60 kg CO_2_ per tree, on average. The nationwide flower TC in each spring was calculated to be 1,900±920 tonnes of CO_2_, equivalent to the yearly carbon capture of 176 hectares of mature pine trees and carbon emissions from 0.24 million car operations each day. As compounds from cherry blossoms can be used extensively for pharmaceutical and cosmetic products, harvesting can be cost effective. Yet, its environmental costs and disposal after component extraction need to be considered altogether in a more complete life cycle analysis of diverting this product from landfill or decomposition.

## Background

One of the major challenges for recent generations of humanity has been global warming and the consequent climate change. Humans, collectively, have tried to adopt solutions to it in two major ways: reducing and quenching carbon dioxide (CO_2_). The first is to cut down the production of CO_2_ through alternatives to burning fossil fuels as energy source. The other is to capture CO_2_ existing in the atmosphere and fixing it to immobile carbon. Technologies like the Climeworks AG facility in Switzerland aim to employ ‘Direct Air Capture (DAC)’ whilst expending some energy to install and pump the captured gas into underground storage (Climeworks.com)^22^. However, plants appear to be more effective in terms of quenching. Mature trees planted in an area of 0.9 billion hectares can reduce about two-thirds of carbon in the atmosphere accumulated since the Industrial Revolution without energy cost (Crowther Lab of ETH Zurich, 2019)^1^.

Flowers are reproductive organs of plants. Unlike stems, leaves and roots, they are temporary in flowering trees. They are also made from atmospheric CO_2_. They are transient in nature and are quickly decomposed into organic molecules of carbon, nitrogen and others, ultimately releasing CO_2_. Thus, harvesting flowers, instead of natural or artificial decomposing, can have the effect of CO_2_ quenching as it prevents some of the CO_2_ from re-entering into the air.

Among all flowering trees, street and garden trees are the most accessible and controllable. The most abundant street flowering tree is cherry blossom in South Korea, consisting 28% of all street trees in Korea as of 2019 (Korea Forest Service, 2020)^2^. Their synchronised blooming in similar climatic regions for a short period of time make the amount of floral waste substantial on streets in spring. Aesthetically pleasing scenery turns into environmental nuisance. It can be hypothesised that harvesting flowers on trees or fallen flowers not only eliminates nuisance but also prevents CO_2_ from re-entering the atmosphere.

This research aims to evaluate the feasibility of reducing atmospheric carbon dioxide through flower harvesting of cherry blossoms. This study estimated the amount of carbon and CO_2_ that can possibly be reduced by flower harvesting and discussed profitable applications of harvested flowers that can promote industrial-scale flower harvesting and waste treatment.

## Methods

### Specimen collection

Samples for measuring carbon contents in flowers were collected in four different locations in the southern part of South Korea (Table 1). For samples g1-6, apical branches 7 - 30 cm long with flowers was cut. They belonged to *Prunus* x *yedoensis* when identified by a set of taxonomic keys on flower and other parts. Keys for flowers used hairy style, cup-shaped calyx tube, and white or pink flowers in bundles of 3-6 flowers as criteria for identifying to species level (Han et al, 2022)^19^. Keys for other parts used gray-brown bark, toothed oval leaves in alternating arrangement with hair on the veins at the back to identify species (Kim, 1998)^3^.

**Table 1.**
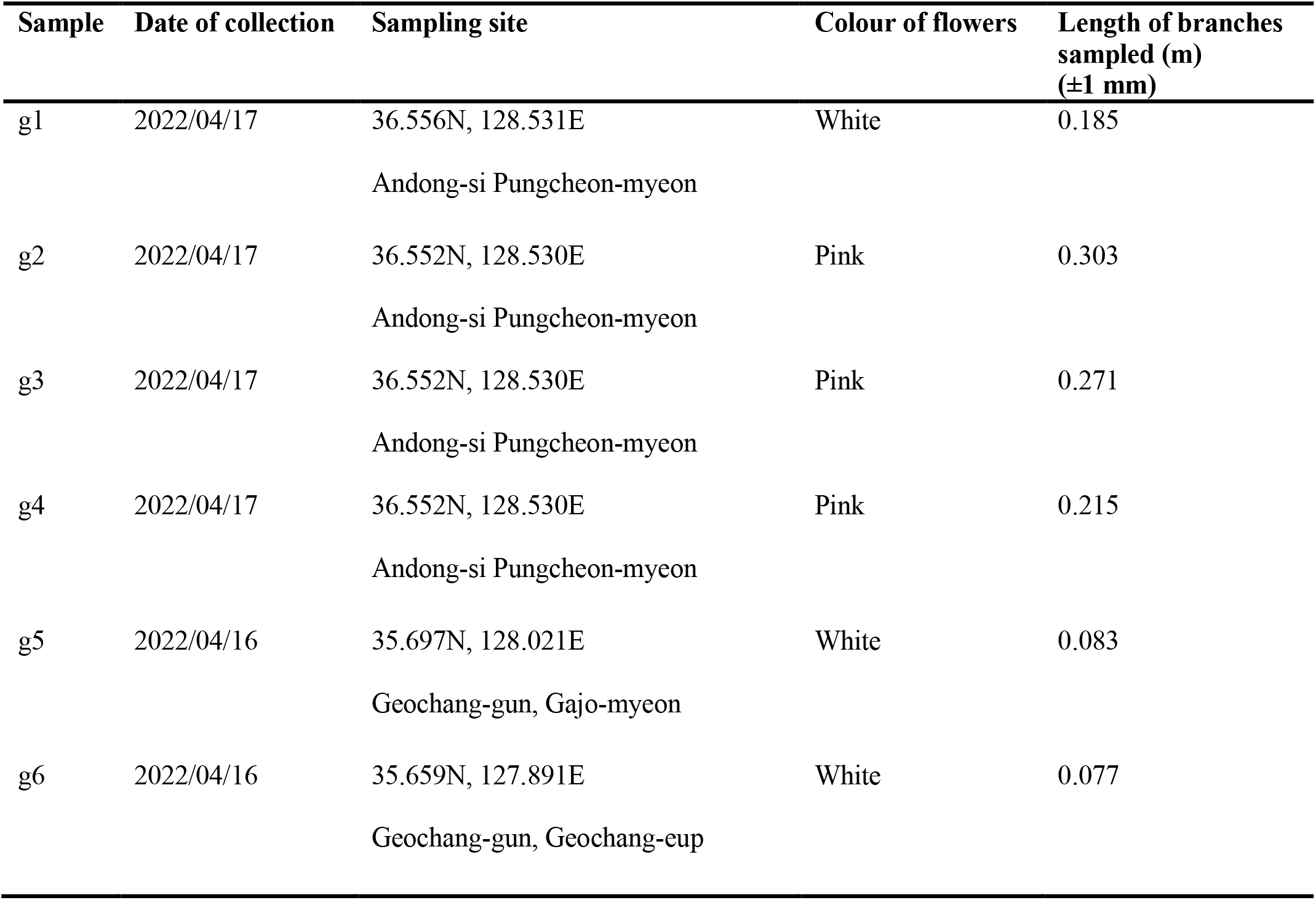
Sample information

### Measurement of total carbon in cherry blossom flowers

Flowers were cut up into receptacles from the stems. Six samples of flowers were dried at 80°C for 48 hours in an oven. Dry weights of each flower sample were measured by precision balance (ED423S, Satorius, Germany) with nominal standard error of measurement ±0.5mg. Dried flower samples were ground with mortar and pestles cleaned with 0.1N HCl. Each sample was put into a labelled vial and sent to a TOC analysis service (Aurora 1030D TOC Analyzer, OI Analytical Corporation, Texas) to determine the average mass of carbon per 50 mg dry weight and the average % carbon content. TC of each ground flower samples were measured in triplicates.

### Calculating the mass of carbon in flowers per tree

#### Amount of flower TC on a branch

The branch lengths of flower samples were measured using a meter ruler with nominal standard error of ±1 mm (Table 1). In addition, the carbon mass in flowers was calculated by multiplying avg. % carbon content by the mass of flowers (dry wt.). The flower TC of each branch was divided by its length; the values were then averaged to produce a mean value of flower TC stored per metre of branch (g/m). It was assumed that flowers were equally distributed along branches with flowers across the tree based on results of a preliminary analysis. The analysis involved counting flowers on apical and subapical branches and determining a linear relationship with branch length or basal area of a branch. Results of the analysis was presented in the Results section.

#### Mean crown diameter of cherry blossom trees on streets of Korea

Twenty regions where abundance of cherry blossom tree was well known were selected for geographically even sampling of street trees. Ten images of tree crowns with contrast to background and discrete crown were randomly selected in each of 20 regions for measurement of their crown diameter. The shortest and the longest crown diameters of each tree were measured on satellite images of tree crown with ‘Distance Measurement’ function in Naver Map application (Naver, Korea; Fig. 1). The mean of all crown diameters for 200 trees was adopted as the average crown diameter of typical cherry blossom trees on streets in South Korea.

**Figure 1.**
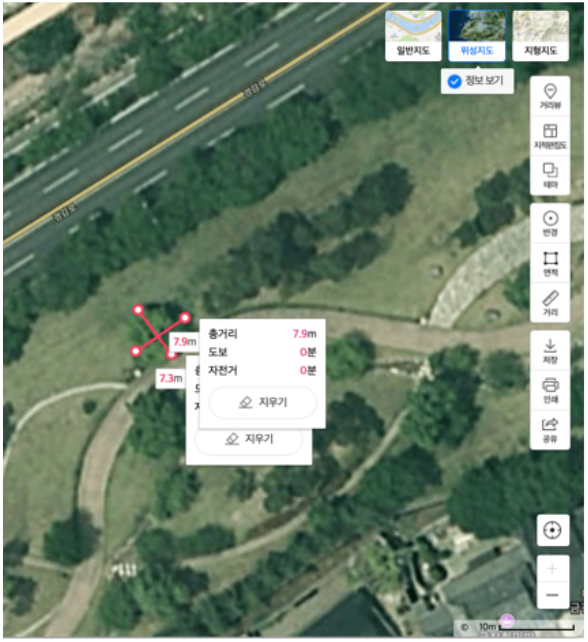
An example of measuring the shortest and the longest crown diameters of a tree.

#### Mean flower TC of cherry blossom trees on streets of Korea

To obtain total flower TC on a tree based on mean flower TC values on a metre of branch, a functional relationship was developed by plotting a regression curve of total length of flowering branches (TLFB) of a tree against its crown diameter. TLFB was determined as apical and subapical branches because cherry blossom trees form their flowers mostly on these branches (see ‘Discussion’ for further details). By applying the mean crown diameter of cherry blossom trees on streets of Korea (described above) as parameter in the function, the mean value of total branch length of a tree on streets of Korea was obtained. The mean value of total flower TC of a tree on streets of Korea was calculated as the product value between the mean value of TLFB and flower TC values on a metre of branch. The process of developing the function converting crown diameter of a tree into TLFB of the tree is described below.

##### Selecting trees for estimating number and length of branches

Trees showing crowns of diameter of 2 - 10 m, circular shape, and interspace from other trees were selected from satellite images on the city of Busan. Among the trees, five trees showing crown diameters approximately 2, 4, 6, 8, or 10 m respectively were randomly selected for non-destructive estimation of branch parameters.

##### Counting the number of branches by branch orders

Branches were identified as their respective orders. Branch order is defined using the framework of Strahler ordering of branches as reference (Fig. 2A). A ‘branch’ was defined as a collection of adjoining segments until the point of separation at the tip (Eloy et al., 2017)^4^. In this investigation, the most apical branch was ranked as the first order and the second most apical as second, and so on.

**Figure 2.**
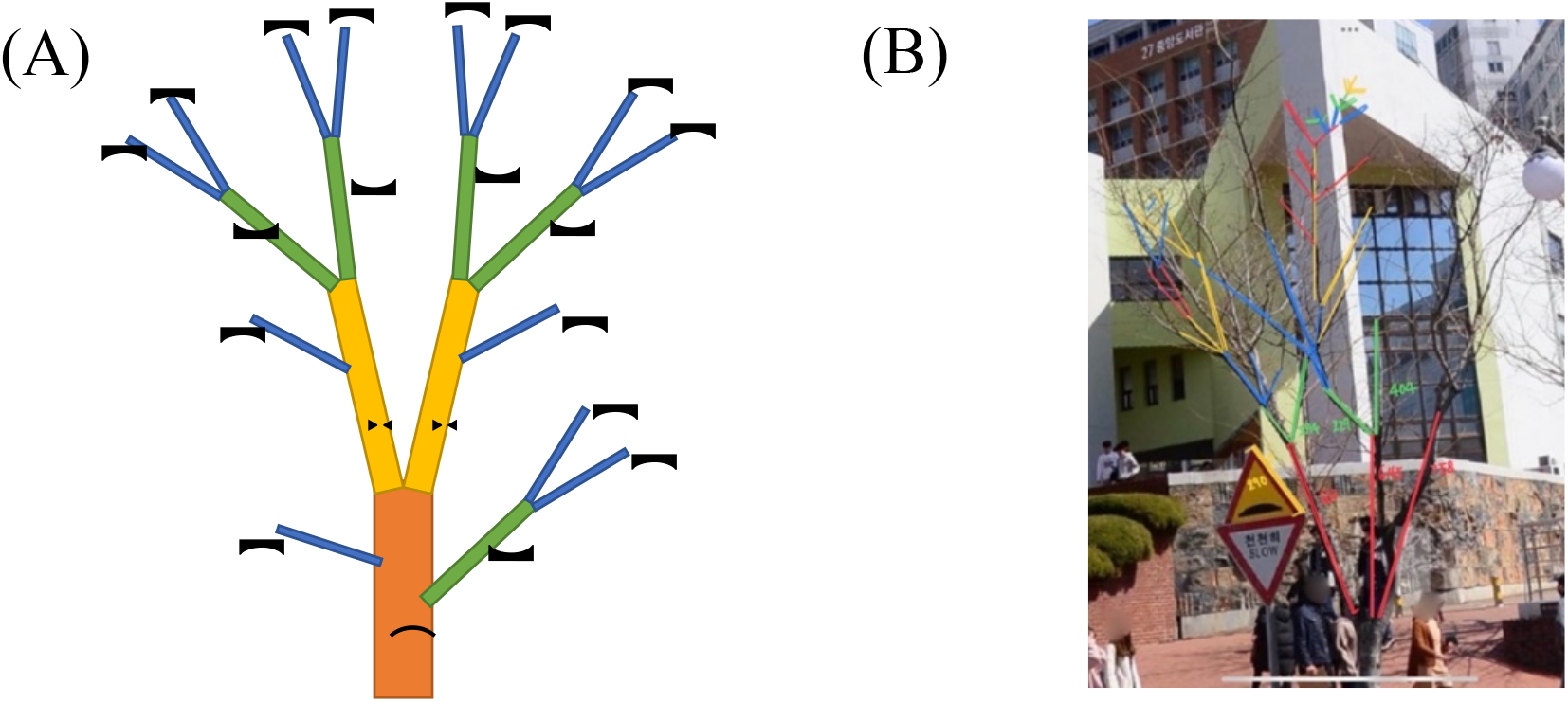
Definition of branch order (A). Schematic illustration for Strahler ordering of branches, (B) An example of assigning orders.

The number of branches of an order was determined by counting the number of branches originated from one branch of the previous order. One of the major branches emerging from the trunk of a tree were selected for counting branches of child orders. The average number of branches at each order were calculated from all child branches originating from the major branches. The total number of branches in a child order was calculated as the product of the mean number of the child order and the mean number of the mother order.

##### Branch lengths at each order

The lengths of apical (i.e., the first order) and subapical (i.e., the second order) branches were measured in app pixel units by the ‘Ruler’ function in Apple Notes app (iOS 15.6) on images of tree branches. Simultaneously, actual lengths of some parts of branches were actually measured non-destructively on site by a laser distance meter (SD-70, Sincon, China) and trigonometric calculation. The ratio of meter unit to app unit was calculated for those parts. The mean length of each order of branches was converted to meter unit using the ratio as the conversion factor.

##### Total length of flowering branches of a tree

The total number and the mean length of branches for the first and the second order was multiplied to give a sum of branch lengths for a sector of the tree crown formed by all branches originating from one basal branch used for branch counting and length measurement. The central angle of the crown sector occupied by child branches of one basal branch was estimated using the satellite image of tree crown and the ‘Angle’ function in Apple Notes. The ratio of the central angle of the sampled sector to 360 degrees were used as conversion factor to extrapolate the total flowering branch length for the whole crown of a tree from the sampled sector.

### Total carbon content in cherry blossom flowers in South Korea

The mean value of total flower TC of a tree on streets of Korea was multiplied by the total number of cherry blossom trees on streets and gardens of South Korea to obtain the nationwide flower TC formed each spring. The tree distribution statistics in Korea were obtained from the Korea Forest Service^2^. The nationwide flower TC was converted into the nationwide CO_2_ equivalent unit by multiplying the ratio of molecular weights of CO_2_ to elemental carbon.

## Results

### Abundance of flowers per unit length of branches order

To extrapolate total amount of flowers on a tree from parameters of its crown and branches, distribution of flowers on branches of different orders was analyzed. For sweet cherry (*Prunus avium* L.), it has been reported that its flower density was proportional to the logarithm of branch length (Jacyna et al, 2021)^17^. For the case of sour cherry (*Prunus cerasus* L.), the number of flowers per basal area of a branch on a tree was reported to be constant (Hrotkó et al, 2008)^18^. In both sweet and sour cherries, flowers formed mostly on apical (branch order 1) and subapical (order 2) branches. In this study, it was checked whether it is branch length or basal area that better determines flower abundance on branches of *P. × yeodoensis*.

For cherry blossom trees in Busan, South Korea, the number of flowers on apical (order 1) and subapical (order 2) branches of various lengths were counted. Flower abundance of the cherry blossom branches were not related to basal areas of subapical branches, and the correlation between flower abundance and basal areas of apical branches was relatively weak (Fig. 3B). However, branch length explained difference of flower abundance by branches with high levels of proportionality for both apical and subapical branches (*R*^2^ > 0.7, Fig. 3A). When both order 1 and order 2 branches were combined, flower abundance was best fitted to a linear equation with the highest *R*^2^ (0.85). This result implied that cherry blossom flowers were distributed evenly along an orders 1 and 2 branches. Therefore, we could estimate flower abundance of a branch by its length using the equation of *N* = 0.53*L* + 3.17, where *L* stood for branch length (cm) and *N* for flower abundance.

**Figure 3.**
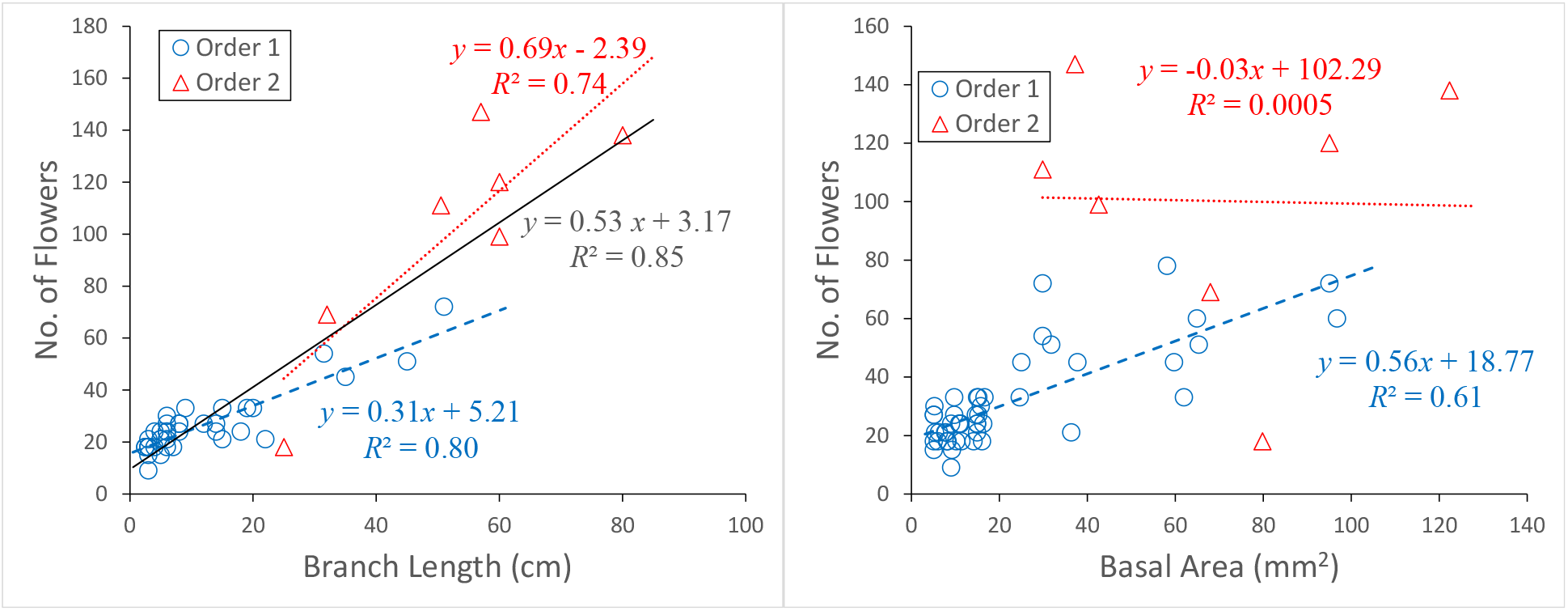
Relationship between length and basal area of apical and subapical branches of a cherry blossom tree.

### Total carbon content in unit mass of flowers

For the six flower samples (g1-g6, Table 1), TC was analyzed by TOC analyser. Per dry weight of flowers, 41.5% - 44.8% of the mass was due to carbon elements out of 367±174 g of flower dry weights (Table 2). These values are similar to the carbon content (45.01±5.23%) of reproductive organ measured for 142 plants sampled globally (Ma et al, 2018)^20^, implying that carbon content of cherry blossom flowers varies by habitat condition to some extent but within the range of that of typical plants. The average percentage carbon content per dry weight of flowers was concluded to be 43.45±2.03%.

**Table 2.** Mass, carbon content and length of samples used to calculate flower carbon on unit length of branch

| Sample | Avg. mass per 50<br>mg dry weight of<br>flowers (mgC) | Avg. %<br>carbon | Mass of<br>flowers<br>(mg) | Flower<br>carbon<br>mass (gC) | Branch<br>length (m) | Flower carbon<br>per metre of<br>branch (gC/m) |
| --- | --- | --- | --- | --- | --- | --- |
| g1 | 22.3±0.75 | 44.6±1.50 | 393 | 0.18 | 0.185 | 0.947 |
| g2 | 21.1±0.70 | 42.2±1.40 | 530 | 0.22 | 0.303 | 0.739 |
| g3 | 21.6±0.79 | 43.3±1.57 | 547 | 0.24 | 0.271 | 0.873 |
| g4 | 22.2±0.74 | 44.4±1.48 | 420 | 0.19 | 0.215 | 0.865 |
| g5 | 22.4±0.83 | 44.8±1.66 | 162 | 0.07 | 0.083 | 0.875 |
| g6 | 20.7±0.65 | 41.5±1.31 | 151 | 0.06 | 0.077 | 0.809 |

### Flower carbon content on unit length of branches

To estimate carbon mass of flowers on the sample branches, dry weights of the flower samples were multiplied by % carbon value for each sample (Table 2). The sample branches were estimated to have 0.06 – 0.24 grams of carbon. By multiplying the length of each sample branch to flower carbon mass of each branch, flower carbon mass per metre of sample branch was obtained (Table 2). On average, the sample branches had 0.851±0.070 gC per metre of branch, and this value was used as the estimate of flower TC per metre of branches of cherry blossom trees in Korea in further analysis.

### Total carbon content in unit length of branches

Since we have the estimate of mean flower TC per metre of branches of cherry blossom trees in Korea, TC in all flowers of a tree could be calculated by estimating total length of all flowering branches of a tree (Table 3).

**Table 3.** Total branch length of a tree: considering the order of branches and the angle occupied by the sampled crown sector

| Tree | Crown diameter (m) | Order | Total number of branches for each order in the sampled sector | Mean length of branches at each order in the sampled sector (m) | Total branch lengths at each order in the sampled sector (m) | Central angle for sampled crown sector | **Total length of flowering branches for whole crown (m)** |
| --- | --- | --- | --- | --- | --- | --- | --- |
| #1 | 2.50 | 1st | 53 | 0.31 | 16.6 | 100° | **108** |
|  |  | 2nd | 16 | 0.85 | 13.5 |  |  |
| #2 | 4.55 | 1st | 207 | 0.15 | 31.2 | 97° | **161** |
|  |  | 2nd | 35 | 0.36 | 12.3 |  |  |
| #3 | 6.00 | 1st | 540 | 0.11 | 57.5 | 108° | **314** |
|  |  | 2nd | 180 | 0.21 | 37.0 |  |  |
| #4 | 8.00 | 1st | 76 | 0.17 | 19.3 | 48° | **413** |
|  |  | 2nd | 52 | 0.57 | 35.8 |  |  |
| #5 | 9.65 | 1st | 631 | 0.25 | 156.1 | 75° | **946** |
|  |  | 2nd | 100 | 0.41 | 40.9 |  |  |

#### Total length of flowering branches in a tree

As a way of summing up lengths of all flowering branches, we estimated an average number of branches and mean branch length at each order. The five trees used as sample cases of trees with different crown sizes showed orders of branches up to 4 – 8. Since most flowers are formed on apical or subapical branches, lengths of the first and the second order were summed. The total number of apical branches reached 631 for the sampled sector of the largest tree. The mean branch length of apical branches was 89±41 cm and that of subapical branches was 225±141 cm, indicating subapical branches are longer by 2.5-fold. The total lengths of branches at the first and the second order was calculated as the products of mean branch number and mean branch length of each order. The total flowering branch length in each sector of a tree crown was calculated by adding branch lengths of the two orders. The sector-wide branch length was extrapolated to the total branch length of whole crown for a tree based on the central angle of the sampled sector. Central angles of the sampled sectors ranged 48-108°, and the extrapolated TLFB in a tree ranged from 108 m for the smallest to 946 m for the largest tree.

#### Function for converting crown diameter to total branch length of a tree

A linear positive correlation could be observed between the crown diameter (*x*) of a tree and the log of its TLFB (*y*), producing a function of *y* = 47.9 *e*^0.30*x*^ by exponential regression (Fig. 4). The estimates were highly significant since 96.6% of variance in the length could be explained by crown diameter. The function allowed us to obtain TLFB of any cherry blossom tree from its crown diameter between 2.5 – 9.6 m by simply measuring its crown diameter.

**Figure 4.**
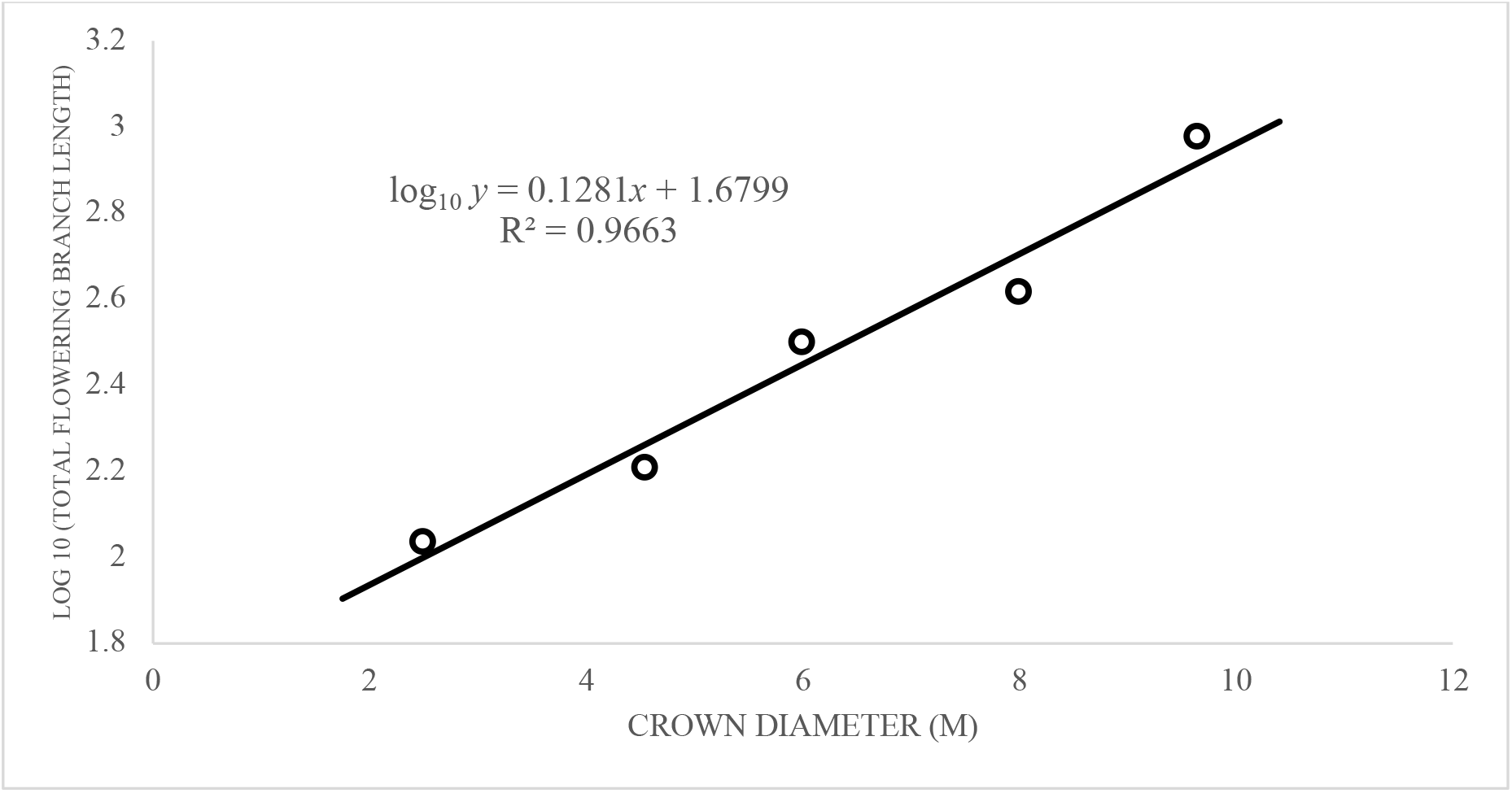
Relationship between the crown diameter and the total length of apical and subapical branches.

#### Estimating flower TC of a tree by its crown diameter

Having an estimate of mean flower TC per metre of branches of cherry blossom trees in Korea (Table 2) and a function to convert crown diameter into TLBF of a tree (Fig. 4), flower TC of a tree could be calculated by multiplying by 0.851 g of flower TC per metre of branches to its TLBF, converted from its crown diameter.

### Nationwide carbon content in cherry blossom flowers in South Korea

The mean crown diameter of cherry blossom street trees across the country was obtained by sampling 200 different trees and taking the average (Table 4). This was then converted into an estimate of total flowering branch length of a typical cherry blossom tree on Korean streets using the conversion function (Fig. 4). The product of TLFB and the mean flower TC in a metre of a branch was multiplied by the number of street trees in the country. This gave the final value indicating TC in all cherry blossoms in South Korea. The estimation of each parameter is presented below.

**Table 4.** Average crown diameter of ten trees selected randomly in each region

| Region | Coordinate of the centre of region | Mean crown diameter (m) | SD (m) |
| --- | --- | --- | --- |
| Hwahweo village, Andong | 36.5389°N, 128.5181°E | 8.98 | 1.92 |
| Andong dam, Andong | 36.5684°N, 128.7294°E | 8.24 | 2.38 |
| Gyeongang-ro, Gyeongju | 35.8365°N, 129.2196°E | 7.40 | 1.88 |
| Daewoo marina, Busan | 35.1602°N, 129.1467°E | 7.96 | 1.69 |
| Kyungsung University, Busan | 35.1398°N, 129.0985°E | 8.67 | 2.35 |
| Hogupo-ro, Incheon | 37.4184° N, 126.7117° E | 6.80 | 0.82 |
| Gwanchok-ro, Nonsan | 36.2011° N, 127.0995° E | 5.19 | 0.76 |
| Sang-dong, Jeong-eup | 35.5605° N, 126.8715° E | 4.95 | 0.80 |
| Jeseok-ro, Gwangju | 35.1208° N, 126.9071° E | 6.13 | 1.59 |
| Daewon-sa, Bosung-gun | 34.9613° N, 127.1330° E | 6.80 | 1.75 |
| Gyeongpo-ro, Gangreung | 37.7839° N, 128.8889° E | 6.57 | 1.05 |
| Parkchunghee-ro, Gumi | 36.1076° N, 128.3529° E | 4.77 | 0.63 |
| Hwagae-ro, Hadong-gun | 35.2268° N, 127.6367° E | 7.93 | 1.13 |
| Yeojuwacheon-ro, Changwon-si | 35.1594° N, 128.6602° E | 10.03 | 1.58 |
| Palgongsan-ro, Daegu | 35.9794° N, 128.6654° E | 5.04 | 0.76 |
| Yeojuiseo-ro, Seoul | 37.5338° N, 126.9202° E | 6.66 | 1.27 |
| Moosim-cheon, Cheongju | 36.3232° N, 127.3452° E | 9.46 | 2.16 |
| Dong-gu, Daegu | 35.8866° N, 128.6353° E | 8.52 | 1.38 |
| Jangseungpo-ro, Geoje | 34.8560° N, 128.7250° E | 7.75 | 1.89 |
| Noksan-ro, Jeju | 33.3903° N, 126.7329° E | 5.24 | 0.96 |

### Crown diameter of typical cherry blossom trees on Korean streets

Ten trees from each of 20 different locations scattered across the country were randomly selected for crown diameter measurement (Table 4). Based on the diameters of the 200 trees, the mean crown diameter of mature cherry blossom trees across the country was estimated as 7.15±1.59 m.

### Total length of flowering branches of a typical cherry blossom tree in Korea

Crown diameter of a typical cherry blossom tree (Table 4) was converted into total length of flowering branches (i.e., orders 1 and 2) using the conversion function (Fig. 4). The mean crown diameter of 7.15±1.59 m corresponded to 394±191 m in TLFB. The product of this length and the mean carbon mass per unit length of branch indicates that flowers of each tree harbor 336±163 g of carbon (Table 5).

**Table 5.** Total carbon content and CO_2_ equivalent (in parenthesis) in cherry blossom street trees in South Korea

| Mean crown diameter (m) | Mean total branch length per tree (m) | Mean carbon mass per meter (gC/m) | Mean total carbon content of flowers per tree (g) | Number of cherry blossom street trees in South Korea (in 2019) | Total carbon content in cherry blossom street trees in South Korea (tonnes per year) |
| --- | --- | --- | --- | --- | --- |
| $7.15 \pm 1.59$ | $394 \pm 191$ | 0.851 | $336 \pm 163$<br>(1231±599) | 1,540,000 | $517 \pm 251$<br>(1,895±922) |

### Estimation of total flower TC of cherry blossom trees across the country

The number of cherry blossom street trees in South Korea was known as 1,540,000 (Korea Forest Service, 2020)^2^. Therefore, approximately 517±251 tonnes of carbon could be quenched by harvesting cherry blossoms on streets of South Korea each year. The estimate corresponded to 1,895±922 tonnes of CO_2_ equivalent.

## Discussion

### Evaluation of the estimated amount of CO_2_ quenched by harvesting flowers

To scale the estimate of CO_2_ that can be quenched by cherry blossom flowers across the country each year, we intended to compare the value to a known value of annual CO_2_ absorption by cherry blossom trees. (Jo and Ahn, 2012)^21^ have produced a function relating the diameter at breast height (*D*_*BH*_) to the amount of CO_2_ absorbed each year (*Y*) for a cherry blossom: ln *Y* = -3.0939 + 1.7702 ln D. To estimate *D*_*BH*_ of a typical cherry blossom tree, a function linking *D*_*BH*_ and the crown diameter was deduced (Fig. 5). Using both functions, the annual CO_2_ absorption for a cherry blossom tree of crown diameter 7.15±1.59 m (equivalent to 32.2±8.8 cm of *D*_*BH*_) was estimated to be 21.1±9.1 kg. This value made the estimated CO_2_ in the flowers of a cherry blossom tree, 1.23±0.60 kg, to be 5.8% of the absorption by cherry blossom trees.

**Figure 5.**
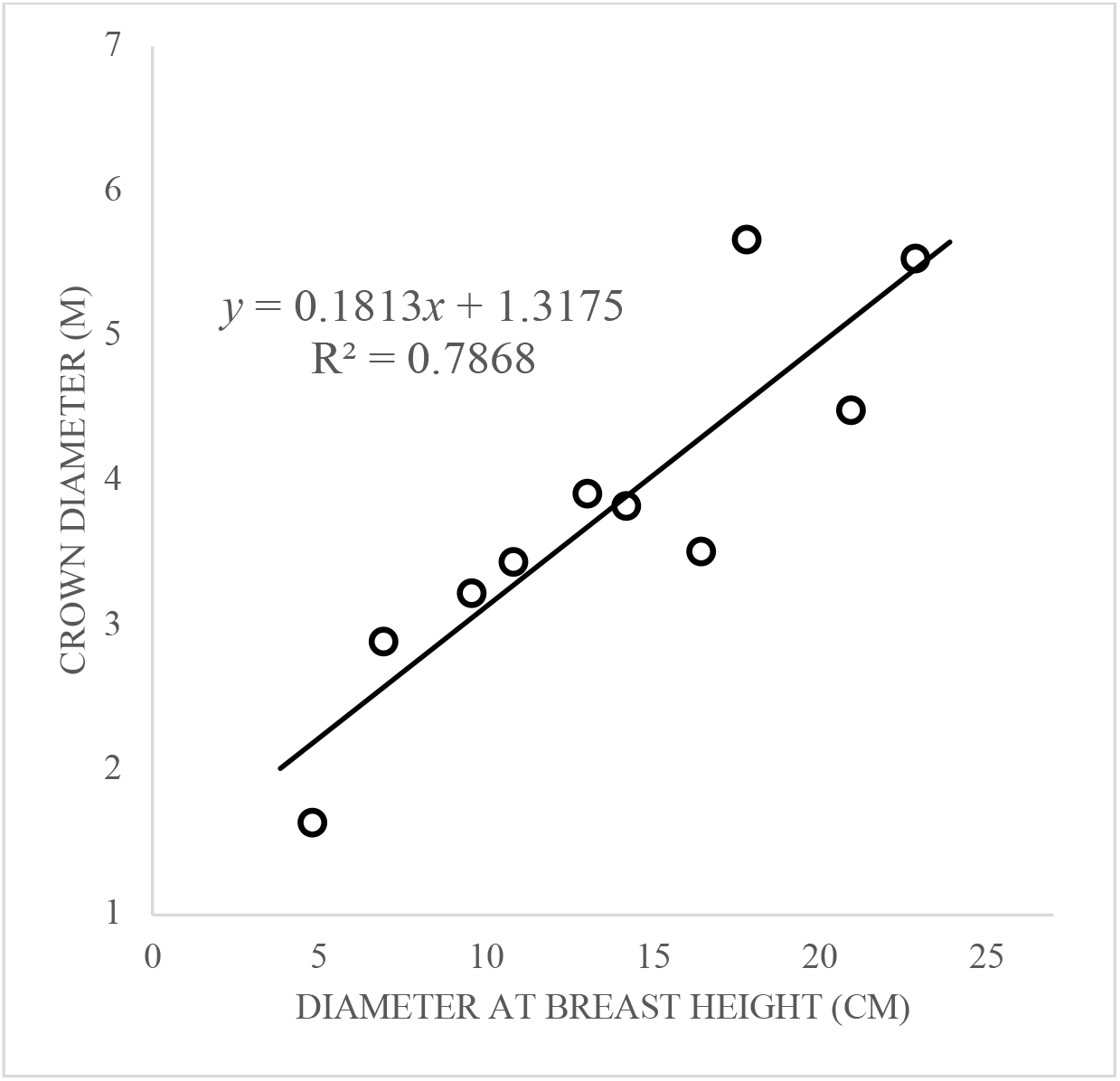
Relationship between *D*_*BH*_ and the crown diameter based on (Jo and Ahn, 2012)^21^

### Equivalence to CO_2_ emissions or other means of reduction

The total CO_2_ emission in Korea in 2019 was estimated to be 586 million tonnes (Korean Statistical Information Service, 2021)^5^. Therefore, 1,900 tonnes of CO_2_ contained in cherry blossoms account for merely 0.00032% of the total annual CO_2_ emission. However, the amount of reduction by harvesting them appears sizeable to other ways of carbon reduction, or emission rates by each well-known source.

To tackle the issue of excessive CO_2_ production, the government has been implementing two major methods: planting trees and replacing fossil fuels with renewable energy sources. As for the former, one hectare of 30-year-old *Pinus densiflora* absorbs 10.8 tonnes of CO_2_ (Korea Forest Service, 2019)^6^, making harvesting the blossoms nationwide each year equivalent to planting 176 hectares of pine trees, an area similar to 44 Wembley stadiums. Employing solar panels as one of the renewable energy sources, 304 tonnes of CO_2_ emission could be decreased (if 1kW per 16.5 m^2^) in one hectare of land per year (Korea Energy News, 2018)^7^. Still, harvesting the blossoms would add 6 hectares of solar panels to existing 6120 hectares of panels in the country since 2015 (Hankyung Economy, 2020)^8^. Considering the cost of installing the panels, estimated to be 5.1 million pounds or 5.8 million USD for 6 hectares in 2018 (Renewable Energy Followers, 2021)^9^, collecting the flowers would be more cost-friendly while reducing carbon dioxide.

In addition, approximately 0.21 kg of CO_2_ is emitted (The Seoul Institute, 2017)^10^ when an internal combustion engine car travels for a kilometre while 1.23±0.60 kg of CO_2_ equivalent is stored in cherry blossom flowers in a tree. This means harvesting the blossoms of one tree has a similar effect of offsetting CO_2_ emission when a car travels for 5.9±2.9 km. A typical car in Korea runs for 37.9 km every day on average (Korea Transportation Safety Authority, 2020)^11^, and emits 8.0 kg of CO_2_ daily. Therefore, harvesting all cherry blossoms on streets across the country once in a year compensate the CO_2_ cost for a daily operation of 0.24 million cars.

### Consequential impacts on the environment by harvesting cherry blossoms

Decomposition is an essential process in nature to circulate the finite nutrients on Earth. Like other parts of plants, flowers are also decomposed by microorganisms in the soil, producing CO_2_ and returning crucial elements such as nitrogen and phosphorous back into the soil for the roots to absorb. When the flowers are from wild cherry blossom trees, removing them from the environment by harvesting would interfere with the nutrient cycle by preventing their re-entry into the soil. However, most leaves or flowers fallen from street trees in Korea often do not enter the soil; instead, they are surrounded by concrete pedestrian ways and asphalt roads. The small area directly underneath the tree, less than 1 m^2^, of exposed soil would not be able to withhold all of the fallen flowers. Moreover, the nutrient contents have already been artificially maintained for street trees by applying fertilisers to meet nutritional requirements for a tree to grow, as they are cultivated trees in a sense. The environmental nuisance is a primary issue after the majority of flowers fall. Accumulated flower petals can cause blockages in the sewage system or inconveniences on roadways and walkways. They are then collected by public cleaners and burned away as garbage, releasing more CO_2_. Although there would be emission due to the transport of the collected flowers, it would not be as significant as the emission by combusting; it would be more environmentally friendly to reuse the petals after harvesting rather than burning them.

### Use of harvested cherry blossom flowers

The authors suggest that the carbon-reducing effect of harvesting cherry blossom flowers is both feasible and substantial as indicated by this small investigation. The question on cost-effective ways of disposing or using the blossoms after harvesting, with minimum carbon emission, therefore arises. In East Asia, there is a long history of using parts of the cherry blossom tree, including the flowers, as herbal medicine or bioactive compounds. Therefore, industrial or commercial value can be added by using the harvested flowers as substances for commercial products, which will motivate flower harvesting and compensate for the costs of harvesting. Nutritional use of flowers can be a traditional example; fresh flowers may have values as animal feed for both terrestrial and aquatic animals, replacing perishable feed ingredients produced by agriculture in spring season. Dried flowers may provide fibers or biomass components for animal feed. The recent development of making cherry flower waste-derived activated carbon with self-doped nitrogen as a supercapacitor and sodium-ion battery could be a cutting-edge industrial application of the harvested flowers (Bhattarai et al, 2022)^12^. Higher commercial values can be also found for harvested cherry blossom flowers as described below.

### Pharmaceutical and cosmetic use of cherry blossom extracts

The extracts from cherry blossoms, particularly the species *P*. × *yedeoensis*, were found out to have proven effects of skin protection against solar UV (sUV) light. In particular, the protein expression of an enzyme crucial for collagen decomposition due to sUV exposure, metalloproteinase-1 (MMP-1), was reduced in the presence of non-enzymatic cherry blossom petal extract (Jung et al., 2021)^13^. This indicates that the extract slows down the degradation of collagen, critical for skin elasticity, implying the role of the blossoms as an anti-wrinkle agent. The idea of using the blossom extract as an antiphotoaging agent was reinforced by the research that demonstrated its ability to protect human keratinocytes from ultraviolet B radiation-induced oxidative stress and apoptosis by regulating critical proteins and pathways involved in the process (Wang et al., 2019)^14^. Furthermore, 0.5-2.0% cherry blossom extract has the effect of reducing NO production during skin inflammation, preventing DNA damage and mutations (Zhang et al., 2014)^15^. Therefore, the harvested cherry blossom flowers can be exploited in skincare products as anti-wrinkle, anti-oxidative and pharmaceutical anti-inflammatory agents that would also be able to produce economic benefits.

### Decomposition in a controlled manner

As mentioned above, much of the cherry blossoms currently collected by street cleaners are burnt in furnaces. An alternative route for the cherry blossoms if not harvested could be discharged into ecosystems through water drainage or trashed as other wet trashes in municipal compost heaps. However, if harvested intentionally for commercial purposes, its decomposition can be controlled in human-maintained facilities. Not all of carbon in the blossoms can be preserved. Yet, carbon emission can be better processed and thus reduced than the case of natural decomposition. Nowadays, biofuel production by biomass decomposition is a common practice (Yuan, 2008)^16^, so that flower decomposition can be coupled to production of renewable fuel partially replacing the use of fossil fuels. Materials after controlled decomposition could be considered as mineral compounds, and it can be used as fertilisers for plant growth. Therefore, flower harvesting can replace some costs for fertilisers. Although there cannot be a waste-free process, results of this study encourage further studies on the ways to utilise the flowers, and the analysis of the cost and benefits entailed by flower harvesting using a more complete life cycle analysis.

## Conclusion

In conclusion, harvesting cherry blossom flowers reduces a substantial amount of atmospheric carbon. 1.9 thousand tonnes of CO_2_ emission, which corresponds to 0.00032% of the total annual CO_2_ emission by all industrial and natural emission processes in Korea, can be reduced by the harvesting. The amount corresponded to planting 176 hectares of pine trees and 0.24 million car operations for a day. Because their extracts can be used extensively for cosmetic products, motivation could exist to harvest them commercially. The environmental costs for harvesting and disposal after component extraction need to be considered as well to implement this suggestion in the real world. This waste product of cherry trees, fallen onto the streets could become an alternative to other virgin materials, e.g., for animal feed. Whilst using mineralised carbon from cherry flower as electronic or ion-battery parts has already been suggested. Cherry blossoms are feasible renewable sources captured carbon and the careful collection and use of this waste could reduce the net carbon footprint on a global scale.

